# CspZ FH-binding sites as epitopes promote antibody-mediated Lyme borreliae clearance

**DOI:** 10.1101/2022.05.24.493343

**Authors:** Yi-Lin Chen, Ashley L. Marcinkiewicz, Tristan A. Nowak, Rakhi Tyagi Kundu, Zhuyun Liu, Ulrich Strych, Maria-Elena Bottazzi, Wen-Hsiang Chen, Yi-Pin Lin

## Abstract

Transmitted by ticks, the bacterium *Borrelia burgdorferi* sensu lato (*Bb*sl) is the causative agent of Lyme disease (LD), the most common vector-borne disease in the Northern hemisphere. No effective vaccines are currently available. *Bb*sl produces the CspZ protein that binds to the complement inhibitor, Factor H (FH), promoting evasion of the host complement system. We previously showed that while vaccination with CspZ did not protect mice from *Bb* infection, mice can be protected after immunization with CspZ-Y207A/Y211A (CspZ-YA), a CspZ mutant protein without FH-binding activity. To further study the mechanism of this protection, herein we evaluated both poly-and monoclonal antibodies recognizing CspZ FH-binding or non-FH-binding sites. We found that the anti-CspZ antibodies that recognize the FH-binding sites (i.e., block FH-binding activity) more efficiently eliminate *Bb*sl *in vitro* than those that bind to the non-FH-binding sites, and passive inoculation with anti-FH binding site antibodies eradicated *Bb*sl *in vivo*. Antibodies against non-FH-binding sites did not have the same effect. These results emphasize the importance of CspZ FH-binding sites in triggering a protective antibody response against *Bb*sl in future LD vaccines.

## INTRODUCTION

Lyme disease (LD) is caused by the spirochete *Borrelia burgdorferi* sensu lato (also known as *Borreliella burgdorferi* sensu lato or Lyme borreliae) transmitted by the bite of infected *Ixodes* ticks. LD is the most common vector-borne disease in the Northern hemisphere, with estimated 476,000 people diagnosed annually in the United States alone (1, 2). Among all spirochete species in the *B. burgdorferi* sensu lato species complex, *B. burgdorferi sensu stricto* (hereafter *B. burgdorferi*) and *B. afzelii* are the major causative species of LD in North America and Eurasia, respectively (2). Following transmission by an infected tick, Lyme borreliae spread from the bite site to various tissues, leading to severe systemic manifestations such as arthritis, carditis, and neuroborreliosis. A human LD vaccine (LYMErix) targeting a spirochete protein, OspA, was once available (but withdrawn from market later), and a second generation of vaccine targeting the same protein is under clinical trial (3-7) However, this protein is not produced after Lyme borreliae invade humans, leading to the difficulty in maintain effective titers of antibodies without constant boosters. Such challenges trigger the need of identifying the new target that are produced in hosts during LD infection as the candidate of LD vaccines.

To survive in a host, Lyme borreliae need to evade multiple host immune responses, while establishing infection and throughout dissemination to distal tissues (8, 9). Part of the immune response in the bloodstream is complement system, which is composed of numerous serum proteins activated through three pathways that recombine to form different protein complexes to ultimately kill pathogens (10-12). Complement regulators are produced to inhibit this cascade to avoid damage to host cells (10, 13). For example, Factor H (FH) and FH-like protein 1 (FHL-1, a truncated form of FH) bind to the C3b in C3 and C5 convertases, leading to the degradation of C3b and inactivating all downstream processes (14). Lyme borreliae produce multiple outer surface proteins that recruit these complement regulators to its surface and promote the degradation of complement proteins upon binding, ultimately facilitating serum resistance and bloodstream survival of the spirochetes (15-21). These spirochete proteins include five distinct FH-binding proteins (collectively known as <u>C</u>omplement <u>R</u>egulator <u>A</u>cquiring <u>S</u>urface <u>P</u>roteins (CRASPs)), including CspZ (CRASP-2) (22, 23). The *cspZ* gene is expressed only when spirochetes reside in vertebrate hosts but not in ticks (24), reflecting the induction of this gene in spirochetes under host environmental cues (e.g., blood and dialysis membrane chambers) (25-27). Additionally, when studied under blood treatment to overcome low expression during *in vitro* culturing, a *cspZ*-deficient *B. burgdorferi* mutant colonizes mouse tissues at reduced levels compared to the wild-type strain (27). These results suggest a role of CspZ to promote efficient spirochete dissemination. Moreover, *cspZ* is highly conserved among Lyme borreliae strains (>80% sequence identity) and carried in all *B. burgdorferi* and *B. afzelii* strains isolated from human patients with systemic and more severe manifestations (e.g., arthritis and neuroborreliosis) (28-30). In addition, all LD patients in an anecdotal study develop antibodies that recognize CspZ (29). These observations raise the possibility of targeting CspZ as a human LD vaccine antigen. However, we and others reported that vaccination with CspZ does not protect mice from *B. burgdorferi* infection (29, 31-33). Rather, immunization with CspZ-Y207A/Y211A (CspZ-YA), a mutant CspZ protein that does not bind to FH, prevents spirochete colonization and LD-associated manifestations in mice (32, 33). These findings lead to the question, ‘*Do antibodies that recognize the FH-binding sites of CspZ confer Lyme borreliae clearance?*’ In this study, we separated the antibodies that selectively recognize CspZ FH-binding or non-FH-binding sites. We then examined each of these antibodies for their ability to eradicate Lyme borreliae and prevent spirochete colonization. The resulting data ultimately facilitated the understanding of the protective epitopes and disease-preventing mechanisms of the CspZ-YA vaccine.

## RESULTS

### CspZ-YA immunization triggered IgGs that recognize both FH and non-FH-binding sites of CspZ

To obtain antibodies that recognize CspZ-YA, sera generated from rabbit immunized with this recombinant protein was purified using a CspZ-immobilized resin to capture all anti-CspZ-YA IgGs (CspZ-YA IgG (Total)) (lane 1 in **Fig. 1A**). From this pool, we separated the IgGs that recognize the FH binding site (CspZ-YA IgG (FH-binding sites)) from those that recognize non-FH-binding sites of CspZ (CspZ-YA IgG (non-FH-binding sites)) (**Fig. 1B**). After applying CspZ-YA IgGs (Total) to a resin functionalized with a CspZ-FH complex, we collected the unbound fraction containing CspZ-YA IgGs (FH-binding sites) (Lane 2 in **Fig. 1A, 1B**). The bound proteins consisting mainly of CspZ-YA IgG (non-FH-binding sites) were then eluted (lane 3 in **Fig. 1A, 1B**). This fraction also contained another protein with a molecular weight of approximately 120 kDa (lane 3 in **Fig. 1A**), likely FH (lane 5 **Fig. 1A**). The bound fraction was subsequently applied to a protein A column to remove the FH, resulting in purified CspZ-YA IgGs (non-FH-binding sites) (lane 4 in **Fig. 1A and 1B**).

**Figure 1.**
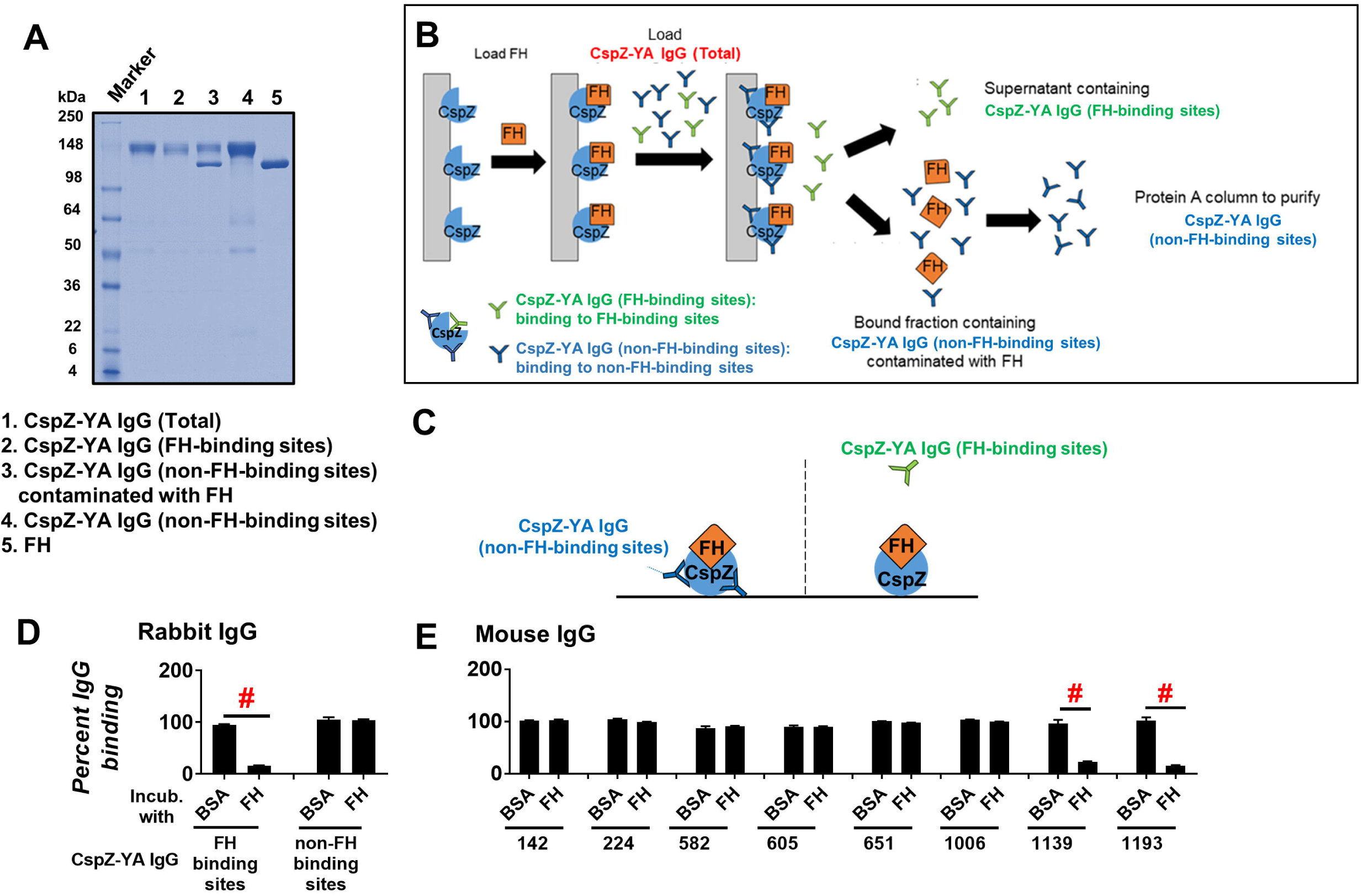
IgGs that recognize both FH and non-FH-binding sites of CspZ can be separately isolated from CspZ-YA-immunized rabbits and mice. **(A)** The integrity and purity assessment of CspZ-YA IgGs using SDS-PAGE. **(B)** A Schematic diagram showing the purification process to retrieve CspZ-YA IgG (FH-binding sites) and CspZ-YA IgG (non-FH-binding sites). CspZ-conjugated resin was incubated with FH followed by loading CspZ-YA IgG (Total). The bound and unbound fractions contain CspZ-YA IgG (non-FH-binding sites) and CspZ-YA IgG (FH-binding sites) IgG, respectively. After eluting the bound fraction, the CspZ-YA IgG (non-FH-binding sites) was further purified using protein A resin to remove contaminated FH. **(C)** A Schematic diagram showing the experimental setup of **(D)** and **(E). (D and E)** One µg of CspZ was coated on ELISA plate wells, which were then incubated with human FH (500nM), BSA (control), or PBS (control, data not shown), followed by the treatment of each of the CspZ-YA IgG samples. These IgGs include **(D)** CspZ-YA rabbit polyclonal IgGs that recognize the FH-binding site of CspZ (FH-binding), or non-FH-binding site (non-FH-binding sites) or **(E)** CspZ-YA mouse monoclonal IgGs (50 nM). The levels of bound CspZ-YA rabbit polyclonal and mouse monoclonal IgG were determined using HRP-conjugated goat anti-rabbit IgG (Sigma-Aldrich) and goat anti-mouse IgG (Sigma-Aldrich), respectively. Data were expressed as the percent IgG binding, derived by normalizing the levels of bound IgG from FH or BSA-treated wells to that in PBS-treated wells. Data shown are the mean ± SEM of the percent IgG binding from three independent experiments. (#) indicates the statistical significance (p < 0.05, Mann-Whitney test) of different levels of percent IgG binding between indicated groups.

To verify the ability of these fractions to recognize (or not) FH-binding sites, we incubated CspZ-coated ELISA plate wells with FH or BSA (negative control), added CspZ-YA IgG (FH-binding sites) or CspZ-YA IgG (non-FH-binding sites) to those wells, and detected the levels of bound antibody **(Fig. 1C)**. We found that CspZ-YA IgG (non-FH-binding sites) bound to both FH- and BSA-treated wells indistinguishably, but the levels of bound CspZ-YA IgG (FH-binding sites) were significantly reduced in FH-treated wells, compared to those from BSA-treated wells **(Fig. 1D)**. These results demonstrate the ability of CspZ-YA IgG (FH-binding sites) to recognize FH-binding sites.

We further investigated whether CspZ-YA immunization triggered the production of IgGs that recognize both FH- and non-FH-binding sites in mice. To generate enough antibodies for this effort and for subsequent experiments, we obtained CspZ-YA mouse monoclonal IgGs from eight individual mouse hybridomas (#142, #224, #582, #605, #651, #1006, #1139, #1193). We then assessed if these antibodies recognize (or not) CspZ FH-binding sites. We found that the binding levels of mAbs #142, #224, 582, #605, #651, and #1006 to FH-treated wells did not significantly differ from those to BSA-treated wells **(Fig. 1E)**. However, antibodies #1139 and #1193 displayed significantly lower levels of binding to FH-treated wells than to BSA-treated wells **(Fig. 1E)**. These results grouped the CspZ-YA mouse monoclonal antibodies into those whose CspZ-binding activity is blocked by FH and those that are not. These findings demonstrate that IgGs that recognize either the FH-or non-FH-binding sites of CspZ can be isolated from CspZ-YA-immunized mice.

### The FH-binding activity of CspZ was selectively blocked by CspZ-YA IgGs that recognize FH-binding sites

Mouse sera after CspZ-YA immunization were documented to block FH-binding to CspZ (32). As the IgGs that recognize either FH-or non-FH-binding sites of CspZ can be isolated after such immunization, we sought to determine the ability of each of those IgGs to prevent FH-binding to CspZ. We thus incubated CspZ-YA IgG (FH-binding sites) or CspZ-YA IgG (non-FH-binding sites) with CspZ-coated ELISA plate wells, added FH to each of those wells, and determined the levels of bound FH **(Fig. 2A)**. CspZ-YA IgGs before fractionation (CspZ-YA IgG (Total)) and irrelevant rabbit IgGs were included as a control. As expected, CspZ-YA IgGs (Total) but not irrelevant rabbit IgG antibodies inhibited FH binding to CspZ **(Fig. 2B, Table S1)**. We found that CspZ-YA IgGs (FH-binding sites) inhibit CspZ-FH binding in a dose-dependent manner, more efficiently than CspZ-YA IgGs (Total) (IC_50_ = 4.2nM). Conversely, CspZ-YA IgGs (non-FH-binding sites) did not inhibit the FH-binding activity of CspZ **(Fig. 2B, Table S1)**. We also determined the capability of each of the mouse monoclonal CspZ-YA IgGs to block FH binding to CspZ. We observed that those IgGs that do not bind to the FH-binding sites of CspZ (#142, #224, 582, #605, #651, #1006, #1139, #1193) did not reduce FH binding to CspZ while those that bound to the FH-binding sites of CspZ (#1139 and #1193) could **(Fig. 2C, see Table S1 for IC**_**50**_). These results indicate that CspZ-YA IgGs that recognize FH-binding sites selectively prevent FH binding to CspZ.

**Figure 2.**
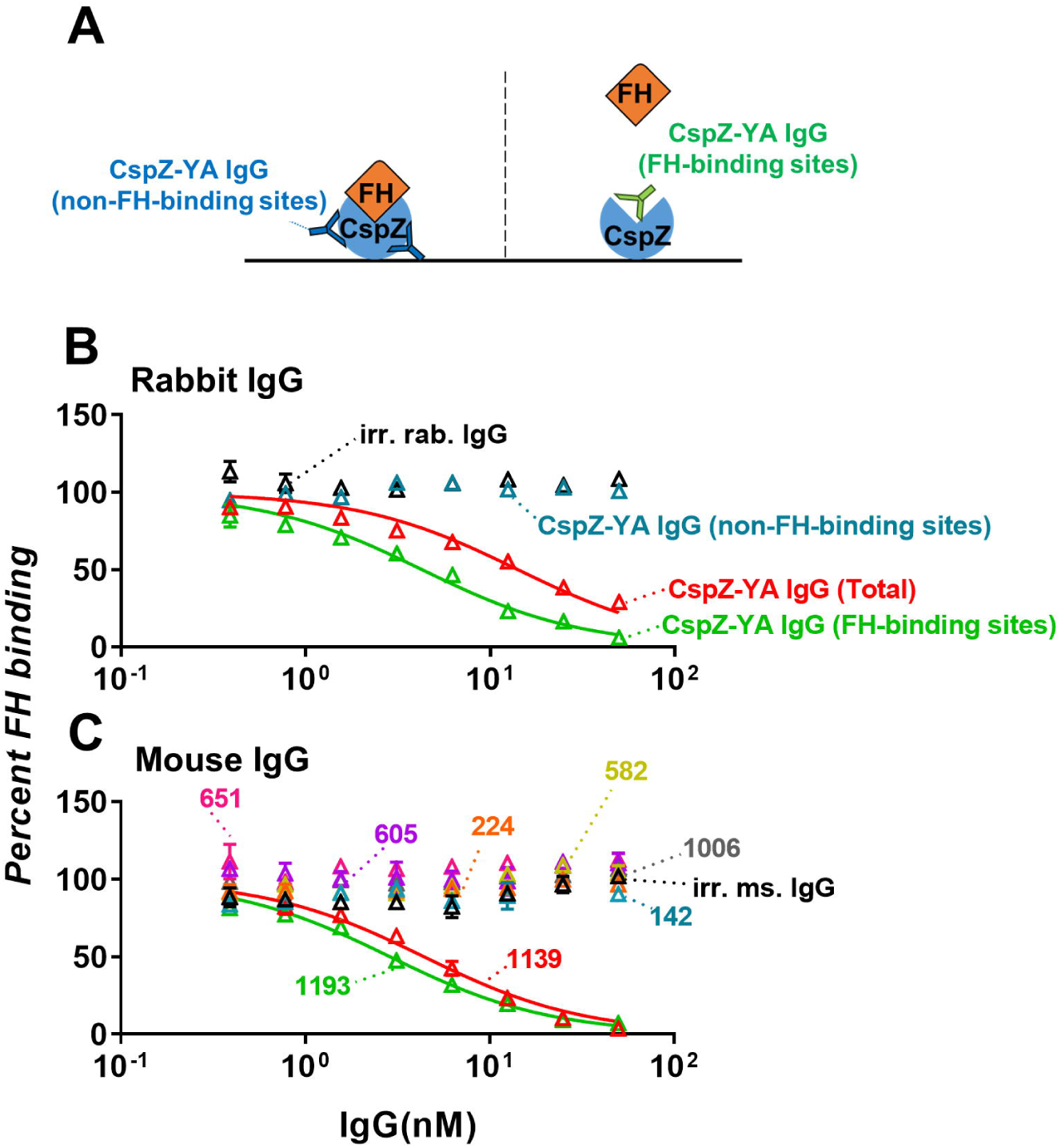
Mouse- and rabbit-derived IgGs that recognize the CspZ FH-binding site selectively block the human FH-binding activity of CspZ. **(A)** Experimental setup. **(B and C)** Each of the CspZ-YA IgGs was added into the CspZ-coated ELISA plate wells. These IgGs include **(B)** CspZ-YA rabbit polyclonal IgG (CspZ-YA IgG (Total)), those IgGs that recognize the FH-binding site of CspZ (CspZ-YA IgG (FH-binding sites)), or non-FH-binding site (CspZ-YA IgG (non-FH-binding sites)), or **(C)** CspZ-YA mouse monoclonal IgGs at indicated concentrations (see Materials and Methods). The wells treated with irrelevant IgGs from rabbits (irr. rab. IgG) and mice (irr. ms. IgG) (50 nM) and PBS (data not shown) were included as a control in the same dose-dependent fashion. Each of those wells was then incubated with human FH, and the levels of bound FH were quantified using sheep anti-human FH and goat anti-sheep HRP IgG as primary and secondary antibodies, respectively. The work was performed on three independent experiments; within each experiment, samples were run in triplicate. Data are expressed as the percent human FH binding, derived by normalizing the levels of bound human FH from IgG-treated wells to that from PBS-treated wells. Data shown are the mean ± SEM of the percent human FH binding from three replicates. Shown is one representative experiment. The concentrations of the IgG to inhibit 50% of human FH bound by CspZ (IC_50_) was obtained from curve-fitting and extrapolation of Panel B and C and shown in Table S1.

### CspZ-YA IgGs that recognize CspZ FH-binding sites robustly killed Lyme borreliae *in vitro*

We have shown that CspZ-YA vaccination triggers borreliacidal antibodies (32), raising the possibility of IgGs that recognize FH- and/or non-FH-binding sites of CspZ to kill Lyme borreliae. We examined this possibility and found that the irrelevant rabbit IgGs do not eliminate *B. burgdorferi* B31-5A4, but CspZ-YA IgGs (Total) killed those spirochetes in a dose-dependent manner (**Fig. 3A**; the 50% borreliacidal activity of each IgG (BA_50_) was 7.4 nM, **Table S2**). We found that CspZ-YA IgGs (non-FH-binding sites) did not efficiently kill *B. burgdorferi* B31-5A4 since the results from this IgG did not allow accurate fitting to obtain a BA_50_ value **(Fig. 3A, Table S2)**. However, CspZ-YA IgGs (FH-binding sites) robustly eradicated those spirochetes, with significantly lower BA_50_ values (BA_50_ = 2.56nM) than CspZ-YA IgGs (Total) **(Fig. 3A, Table S2)**. We also tested the bactericidal activity of each of the CspZ-YA mouse monoclonal IgGs in the same fashion and observed that irrelevant mouse IgG antibodies did not kill *B. burgdorferi* B31-5A4 **(Fig. 3B)**. We found that the IgGs that did not recognize the FH-binding sites of CspZ could be divided into two groups based on their bactericidal activities: Those IgGs that did not have any borreliacidal activities (#224, #582, #1009, **Fig. 3B**) and those that still killed bacteria but were not efficient enough to acquire accurate BA_50_ values (#142, #605, and #651, **Fig. 3B, Table S2**). On the contrary, both IgGs that recognized the FH-binding sites of CspZ (#1139 and #1193, **Fig. 2C**) efficiently killed spirochetes, with #1193 (BA_50_ = 1.23 nM) eradicating bacteria more efficiently than #1139 (BA_50_ = 3.45 nM) **(Fig. 2C, Table S2)**. We further evaluated the ability of the IgGs that robustly killed *B. burgdorferi* B31-5A4 (CspZ-YA IgG (Total), CspZ-YA IgG (FH-binding sites), #1139, and #1193) to eradicate other strains and species of Lyme borreliae, including *B. burgdorferi* 297 and *B. afzelii* VS461. Similar to the results for *B. burgdorferi* B31-5A4, these IgGs eradicated *B. burgdorferi* 297 and *B. afzelii* VS461 in a dose-dependent manner **(Fig. 3C to 3E and Table S2)**. However, CspZ-YA IgGs (FH-binding sites) more robustly killed strain 297 than CspZ-YA IgGs (Total) but showed no significantly different efficiency in killing strain VS461, compared to CspZ-YA IgGs (Total) **(Fig. 3C and 3E and Table S2)**. In addition, #1139 displayed indistinguishable ability from #1193 to eliminate strain 297 but showed more robust killing of strain VS461 **(Fig. 3D and 3F and Table S2)**. These results not only showed the general ability of CspZ-YA IgGs that recognize the CspZ FH-binding sites to efficiently kill Lyme borreliae, but also exhibited that these antibodies varied in their efficiency against different bacterial strains and species.

**Figure 3.**
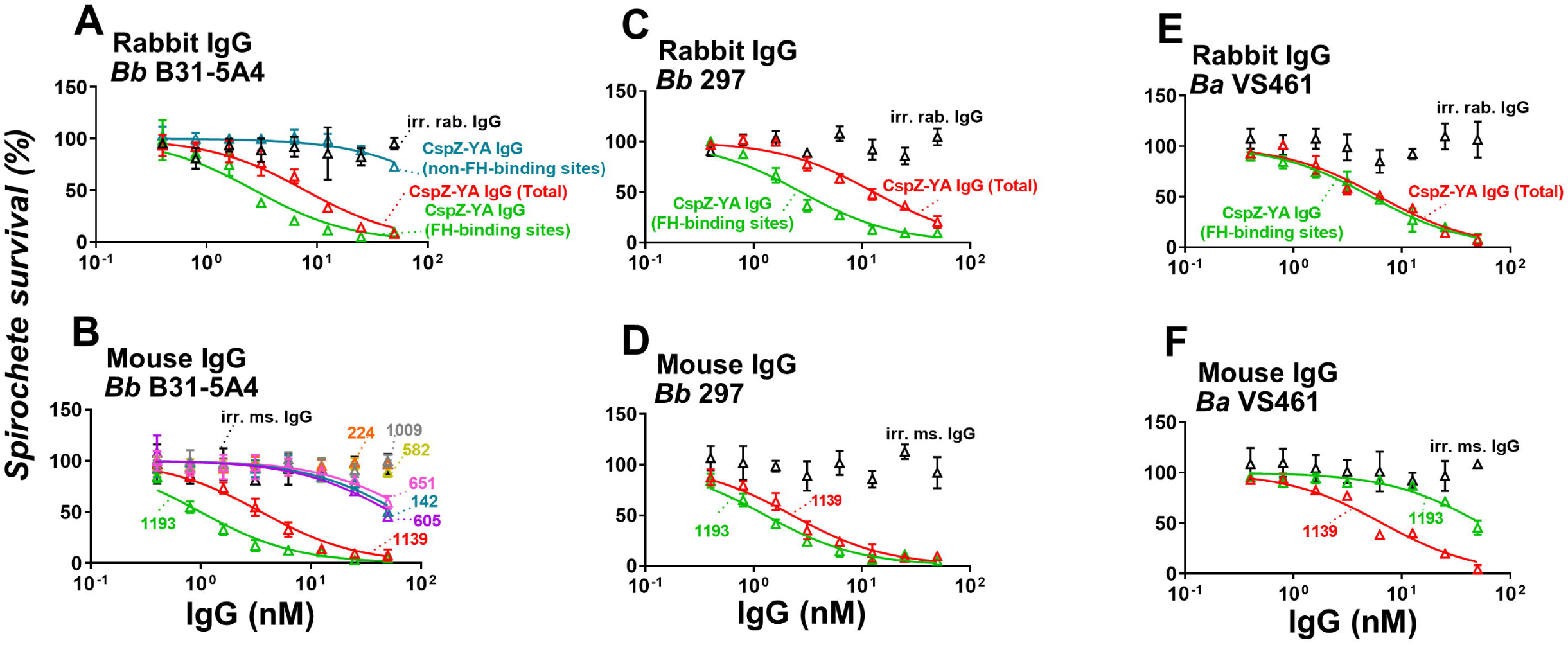
IgGs that recognize the CspZ FH-binding sites more efficiently eliminate different Lyme borreliae species and strains than those that bind to CspZ non-FH-binding sites. Each of the CspZ-YA IgG samples or irrelevant IgG samples from rabbits (irr. rab. IgG) or mice (irr. ms. IgG), or PBS (control, data not shown) were serially diluted as indicated, and mixed with guinea pig complement and **(A and B)** *B. burgdorferi* strains B31-5A4 (*Bb* B31-5A4) or **(C and D)** 297 (*Bb* 297), or **(E and F)** *B. afzelii* strain VS461 (*Ba* VS461) (5 × 10^5^ cells ml^-1^). These IgGs include **(A, C, E)** CspZ-YA IgG (CspZ-YA IgG (Total)), those IgGs that recognize the FH-binding site of CspZ (CspZ-YA IgG (FH-binding sites)), or non-FH-binding site (CspZ-YA IgG (non-FH-binding sites)), or **(B, D, F)** indicated CspZ-YA mouse monoclonal mouse IgGs at indicated concentrations. After incubated for 24 hours, surviving spirochetes were quantified from three fields of view for each sample using dark-field microscopy. The work was performed on three independent experiments. The survival percentage was derived from the proportion of IgG-treated to PBS-treated spirochetes. Data shown are the mean ± SEM of the survival percentage from three replicates. Shown is one representative experiment. The 50% borreliacidal activity of each IgGs (BA_50_), representing the IgG concentrations that effectively killed 50% of spirochetes, was obtained and extrapolated from curve-fitting and shown in Table S2. Data shown are the mean ± SEM of the borreliacidal titers from three experiments.

### Passive immunization with CspZ-YA IgGs that recognize CspZ FH-binding sites reduced the seropositivity and levels of colonization by *B. burgdorferi*

Our previous efforts illustrate that passive inoculation of sera from CspZ-YA vaccination protect mice from Lyme borreliosis (32). This observation led to the hypothesis that the IgGs that recognized FH- and/or non-FH-binding sites of CspZ prevent *B. burgdorferi* infection. To test this hypothesis, we inoculated mice with CspZ-YA IgGs (non-FH-binding sites). Due to insufficient CspZ-YA IgGs (FH-binding sites) recovered after purification for this experiment, mAbs #1139 and 1193 were solely used to represent IgGs that recognize CspZ FH-binding sites. Those mice were then fed on by ticks carrying *B. burgdorferi* B31-5A4 at 1-day post-immunization **(Fig. 4A)**. In addition to uninfected mice, we also included the B31-5A4-infected mice that were inoculated with CspZ-YA IgG (Total), irrelevant rabbit IgG, or irrelevant mouse IgG as a control **(Fig. 4A)**.

**Figure 4.**
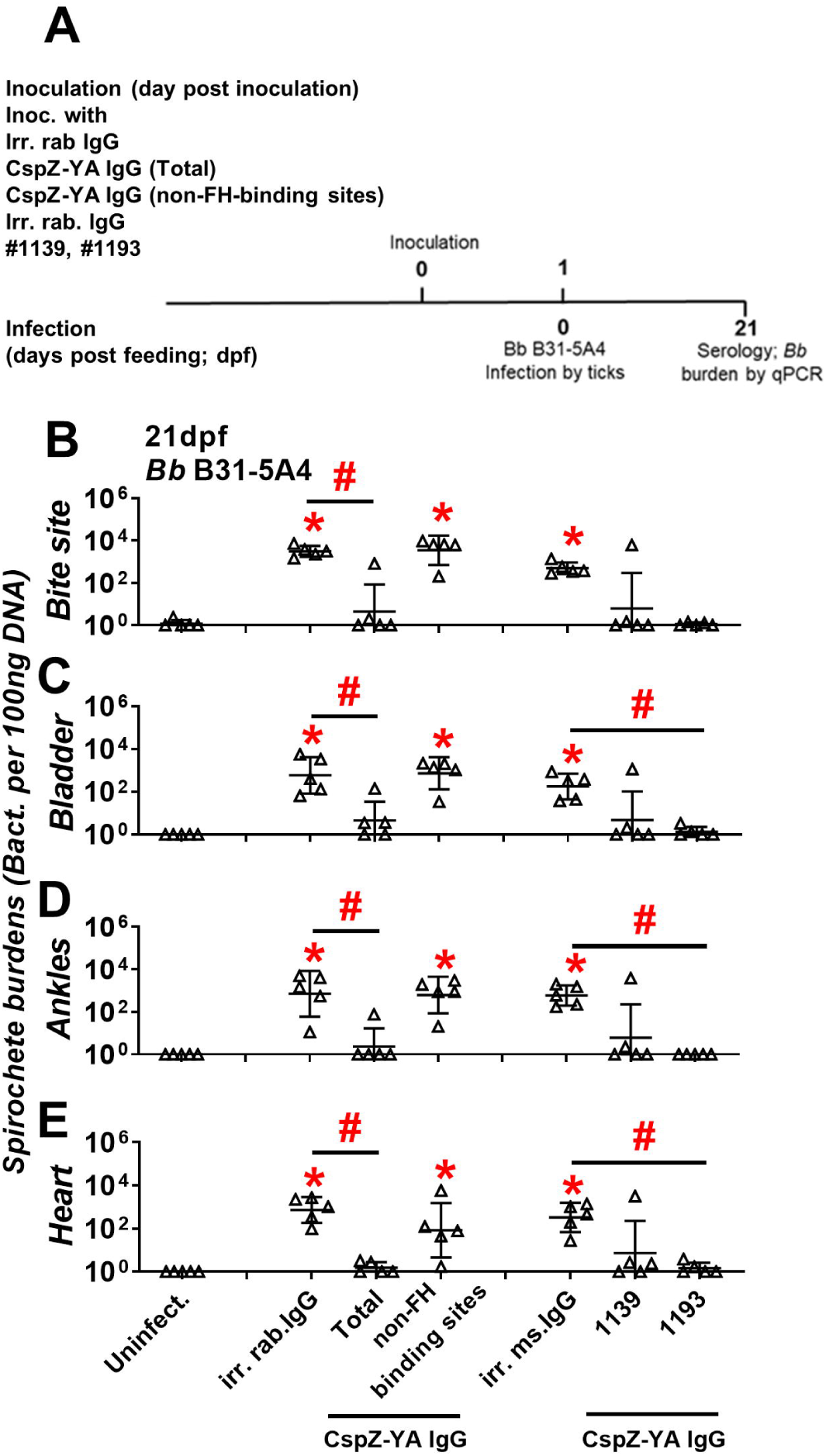
CspZ-YA IgGs that recognize CspZ FH-binding sites selectively prevent *B. burgdorferi* B31-5A4 infection. **(A)** Timeframe of the IgG inoculation and *B. burgdorferi* infection. **(B to E)** C3H/HeN mice were inoculated with irr. IgG from rabbits (irr. rab. IgG) or mice (irr. ms. IgG), or CspZ-YA IgG samples (1 mg/kg, five mice per group). These CspZ-YA IgGs include total CspZ-YA IgG (Total), those IgGs that recognize non-FH-binding site (non-FH-binding sites), or mouse monoclonal IgG #1139 or 1193. At 24 hours after IgG inoculation, these mice were fed on by *I. scapularis* nymphs carrying *B. burgdorferi* B31-5A4 (*Bb* B31-5A4). An additional five mice inoculated with PBS but not fed on by ticks were included as the control (Uninfect.). The tissues were collected from those mice at 21dpf. Spirochete burdens at **(B)** the tick feeding site (“Bite Site”), **(C)** bladder, **(D)** ankles, and **(E)** heart were quantitatively measured at 21 dpf, shown as the number of spirochetes per 100ng total DNA. Data shown are the geometric mean ± geometric standard deviation of the spirochete burdens from five mice per group. Statistical significances (p < 0.05, Kruskal-Wallis test with the two-stage step-up method of Benjamini, Krieger, and Yekutieli) of differences in bacterial burdens relative to (*) uninfected mice or (#) between indicated groups of mice are presented.

We found that after feeding on all groups of mice to full engorgement, ticks did not have significantly different levels of bacterial burdens **(Fig. S1)**, consistent with the finding that CspZ is not produced in ticks (24). At 21-days post tick feeding (dpf), we determined the serology of all mice. The uninfected group was seronegative, whereas all five mice inoculated with irrelevant rabbit and mouse IgG were seropositive **(Fig. S2, Table 1)**. Only one of five mice inoculated with CspZ-YA IgGs (Total) turned seropositive, a significantly lower number than the irrelevant control **(Fig. S2, Table 1)**. Conversely, all five mice injected with CspZ-YA IgGs (non-FH-binding sites) were seropositive, not significantly different from the rabbit IgG control **(Fig. S2, Table 1)**. Notably, only one #1139-inoculated, and none of the #1193-inoculated, mice were seropositive **(Fig. S2, Table 1)**. These results suggest that IgGs that recognize the FH-binding sites of CspZ reduced seroconversion of mice during infection.

**Table 1.** Seropositivity of mice passively immunized with anti-CspZ-YA IgGs followed by the infection of *B. burgdorferi*.

| Inoculated<br>with | PBS | Rabbit IgG |  |  | Mouse IgG |  |  |
| --- | --- | --- | --- | --- | --- | --- | --- |
|  |  | Irr. rab.<br>IgG<sup>c</sup> | CspZ-YA<br>IgG (Total) | CspZ-YA IgG (non-FH-binding<br>sites) | Irr. ms.<br>IgG<sup>d</sup> | 1139 | 1193 |
| Infected<br>with | Uninfect.<sup>b</sup> | *I. scapularis* nymphs carrying *B. burgdorferi* 5A4 |  |  |  |  |  |
| Pos. rate<sup>a</sup> | 0/5 | 5/5<sup>e</sup> | 1/5 | 5/5<sup>e</sup> | 5/5<sup>e</sup> | 1/5 | 0/5 |

<sup>a</sup>Seropositive rate, the number of mice with sera yielding the levels of IgG against C6 peptides greater than the threshold (mean ± three- fold standard deviation of the C6 IgG levels derived from the PBS-inoculated, uninfected mice).
<sup>b</sup>Uninfected mice.
<sup>c</sup>Irrelevant rabbit IgG, anti-green fluorescence protein of rabbit IgG.
<sup>d</sup>Irrelevant mouse IgG, anti-green fluorescence protein of mouse IgG.
<sup>e</sup>Significant greater positive rates compared to the uninfected mice determined by two-tailed Fisher test.

We further determined the levels of spirochete colonization of mouse tissues at 21dpf. Mice inoculated with the irrelevant rabbit IgG control yielded significantly higher bacterial burdens at the bite site, in the bladder, at the ankles, and in the heart than uninfected mice **(Fig. 4B to E)**. Mice inoculated with CspZ-YA IgGs (Total) displayed significantly lower spirochete burdens for all tested tissues than the irrelevant rabbit IgG control **(Fig. 4B to E)**. Conversely, mice injected with CspZ-YA IgG (non-FH-binding sites) showed bacteria burdens indistinguishable from the irrelevant rabbit IgG control, and were significantly higher than uninfected mice **(Fig. 4B to E)**. Additionally, mice inoculated with irrelevant mouse IgG control developed significantly greater bacterial loads in all tissues than uninfected mice **(Fig. 4B to E)**. Except one non-protected mouse in the #1139-inoculated group, all #1139- and #1193-inoculated mice yielded indistinguishable bacterial burdens in all tissues from uninfected mice. Additionally, the #1193-inoculated mice had significantly lower spirochete loads in the distal tissues than the irrelevant mouse IgG-inoculated control mice. These results demonstrate the ability of IgG antibodies that recognize the FH-binding sites of CspZ to significantly reduce *B. burgdorferi* colonization.

## DISCUSSION

The antibody repertoire induced by a specific antigen comprises different populations of antibodies that recognize distinct epitopes on that antigen. Some less abundant antibody fractions are considered “immunologically subdominant” (34). When the efficacious antibody population is immunologically subdominant, host adaptive immune responses may not efficiently eliminate pathogens and/or alleviate manifestations despite overall robust antibody titers (35). Likewise, if the protective epitopes induced by a vaccine antigen were immunologically subdominant that vaccine may be unable to prevent infection (36). Antigen engineering can enhance the abundance of the antibodies that recognize those immunologically subdominant protective epitopes, thereby improving vaccine efficacy (34). One of the strategies for antigen engineering is to remove the structures preventing exposure of the protective epitopes (e.g., surface polysaccharides that mask the protective epitopes of some viral antigens (37, 38)).

The antigen of interest, CspZ, is upregulated immediately after spirochetes infect hosts and binds to host FH (22, 24). Therefore, any protective epitopes in/around the FH-binding site may not be fully exposed to induce sufficient bactericidal antibodies (32, 33). In fact, high titers of antibodies against CspZ are in humans and mice after exposure to Lyme borreliae, but those spirochetes still persist during infection (28, 30, 32, 33). Additionally, we previously reported null protection in mice after immunization with CspZ, but full protection after vaccination with CspZ-YA, a CspZ mutant unable to bind its target ligand (28, 31-33). Thus, the protective epitopes may be immunologically subdominant, and engineering the antigen to remove the FH binding ability may have triggered the production of these protective antibodies. In the current study, we demonstrated the ability of the antibodies that specifically recognize the CspZ FH binding site to eradicate *B. burgdorferi in vitro* and *in vivo*, demonstrating that the protective epitopes were in/around the FH-binding site. Such an engineering of removing the ability to bind the target ligand in enhancing the levels of the protective antibodies has also been applied to other infectious agents, such as Fhbp, a *Neisseria meningitidis* antigen currently used as a human vaccine against meningococcal infection (39-41). However, unlike CspZ-YA, removing the FH-binding activity of Fhbp also magnified overall antibody responses (39, 42, 43). Taken together, our findings provide mechanistic insights into how such a precise antigen engineering would turn immunologically subdominant epitopes to be the major region that can induce efficacious antibodies against infectious agents.

However, an unanswered question is how those antibodies that recognize the mutant FH-binding sites promote spirochete clearance to protect mice from *B. burgdorferi* infection. We showed here that the protection from LD *in vivo* is selectively mediated by the CspZ-YA IgGs that recognize CspZ FH-binding sites. Together with another finding in the current study of antibody-mediated borreliacidal abilities *in vitro* by the IgGs that recognize CspZ FH-binding sites, it is possible that CspZ-YA IgGs-mediated killing is conferred by the Fc regions of IgGs. As that Fc region promotes classical pathway-mediated pathogen lysis and opsonophagocytosis directly or indirectly by the binding of antibodies to leukocytes (44), these mechanisms would play a role in the bacterial clearance caused by CspZ-YA IgGs. We also found that the CspZ-YA IgGs that recognize FH-binding sites blocks FH binding to CspZ, which leads to the possibility that these antibodies eliminate bacteria by preventing the ability of spirochetes to evade alternative pathway-mediated killing (e.g., opsonophagocytosis and pathogen lysis) (10). Delineating the role of each of these mechanisms in promoting the protective antibodies upon CspZ-YA vaccination warrants further investigations.

We found that the CspZ-YA IgGs that recognize FH-binding sites efficiently eliminate spirochetes across different strains and species of Lyme borreliae, but the extent of killing varied among the strains. This result is consistent with the fact that the sequences on the FH-binding interface of CspZ are highly similar among the variants of different strains within the same Lyme borreliae species (>98% identity) but moderately variable among the strains from different spirochete species (∼80% identity) (29, 32). Moreover, CspZ-YA vaccination not only stops spirochete colonization at tissues but also prevents the development of LD-associated manifestations (i.e., arthritis) (30, 32). Although Lyme borreliae dissemination is a prerequisite for the onset of these manifestations, the severity of such disease symptoms has been shown to not necessarily be linked to the spirochete burdens in tissues (45). Therefore, the results from this study showing the absence of the spirochetes in the tissues from mice inoculated with CspZ-YA IgGs that recognize FH-binding sites may not fully address the mechanisms of manifestation prevention by CspZ-YA vaccines. Despite that, our finding does not preclude the possibility of the antibodies that recognize CspZ FH-binding sites killing spirochetes at the initial infection sites and blocking spirochetes from disseminating to distal tissues. In this study, the insufficient polyclonal CspZ-YA IgG (FH-binding sites) from CspZ-YA-immunized rabbits (∼25µg isolated from six rabbits) justifies the use of #1139 and #1193 for the *in vivo* work. However, both #1139 and #1193 may account for only part of the antibody population in the CspZ-YA IgG (FH-binding sites). The fact of the protectivity provided by #1139 and #1193 does not rule out the possibility that the uncovered antibodies population in CspZ-YA IgG (FH-binding sites) have different phenotypes from #1139 and #1193, which is worth further investigation. Further, pre-exposure prophylaxis (PrEP) is a commonly studied strategy for LD prevention. In fact, in addition to vaccines, a yearly administrated monoclonal antibody against OspA, a protein that is required for tick-to-host transmission of spirochetes, is currently in clinical trials (46, 47). Our finding of the bactericidal effect by the antibodies recognizing the CspZ FH binding site illustrates that identifying monoclonal antibodies that bind these epitopes constitutes a promising option for a prophylactic agent against LD. Collectively, this study identified that the protective epitopes of CspZ-YA vaccines are proximal to the FH-binding site, which further elucidates the potential preventive mechanisms of this vaccine candidate. Such findings would eventually inform the strategy of precision antigen engineering in developing vaccines and monoclonal antibody-based prophylaxis against LD and other infectious diseases.

## MATERIALS AND METHODS

### Ethics Statement

All mouse experiments were performed in strict accordance with all provisions of the Animal Welfare Act, the Guide for the Care and Use of Laboratory Animals, and the PHS Policy on Humane Care and Use of Laboratory Animals. The protocol (Docket Number 19-451) was approved by the Institutional Animal Care and Use Agency of Wadsworth Center, New York State Department of Health. All efforts were made to minimize animal suffering.

### Mouse, ticks, and bacterial strains

Three-week-old, female C3H/HeN mice were purchased from Charles River (Wilmington, MA, USA). Although such an age of the mice has not reached sexual maturity, the under development of immune system in this age of mice would allow such mice to be more susceptible to Lyme borreliae infection, increasing the signal to noise ratio of the readout. That will also provide more stringent criteria to define the protectivity. BALB/c C3-deficient mice were from in-house breeding colonies (48) and *Ixodes scapularis* tick larvae were obtained from BEI Resources (Manassas, VA). *Escherichia coli* strain BL21(DE3) and derivatives were grown at 37°C or other appropriate temperatures in Luria-Bertani broth or agar, supplemented with kanamycin (50µg/mL). *Borrelia* strains were grown at 33°C in BSK II complete medium (49), and these strains include *B. burgdorferi* strains B31-5A4 (50), 297 (31, 51), and VS461 (52)(**Table S3**). Cultures of *B. burgdorferi* strain B31-5A4 were tested with PCR to ensure a full plasmid profile before use (53, 54) whereas *B. burgdorferi* strain 297 and *B. afzelii* strain VS461 were maintained as fewer than 10 passages.

### Cloning, expression and purification of CspZ and CspZ-YA

The DNA encoding CspZ and CspZ-YA with a C-terminal TEV cleavage site (ENLYFQG) followed by a hexahistidine tag (his-tag) were codon-optimized based on *E. coli* codon usage preference and synthesized by GenScript (Piscataway, NJ, USA), followed by subcloning into the pET41a using NdeI/XhoI restriction sites. These plasmids were transformed into *E. coli* BL21 (DE3). The recombinant protein expression was induced with 1 mM Isopropyl-β-D-1-thiogalactopyranoside (IPTG). Once expression was confirmed, the clone with the highest expression for each construct was selected to create glycerol seed stocks.

To generate his-tagged CspZ and CspZ-YA, 9 L of Basal Salt Medium (BSM, 5 g/L of K_2_HPO_4_, 3.5 g /L of KH_2_PO_4_, 3.5 g/L of (NH_4_)_2_HPO_4,_ 15 g/L of glucose, 5 g/L of yeast extract, pH 7.2) was prepared and autoclaved. Before inoculation, 9 mL of K-12 Trace Salts Solution, 36 mL of 25% MgSO_4_-7H_2_O, 9 mL of 10% antifoam, 9 mL of 50 mg/mL kanamycin and 90 mL of 15 g/L CaCl2 · 2H2O were added aseptically. 9 L of BSM was then inoculated with a CspZ or CspZ-YA seed culture to a starting OD_600_ of 0.05. The culture was grown at 37°C until OD_600_ reached 0.5-1.0 and the induction phase was initiated. During the induction phase, the culture was induced with 0.2 mM IPTG at 22°C and pH 7.2. After 8 hours of induction, fed-batch feeding (50% v/v glucose) was optionally added at 2 mL/L/h. 30% dissolved oxygen (DO) was maintained throughout fermentation. The cell paste was harvested after 19-hour induction by centrifugation and stored at -80°C until purification. To further purify his-tagged CspZ or CspZ-YA, the cell paste was first thawed and resuspended in 50 mM phosphate buffer with 500 mM NaCl and 20 mM imidazole at pH 7.4 at a ratio of 20 mL buffer per gram and homogenized 3 times on ice at 15,000 psi with an EmulsiFlex-C3 high-pressure homogenizer. The lysed cells were centrifuged and only the supernatant with the soluble proteins was then further filtered with 0.45 µm filters and loaded onto two connecting 5 mL HisTrap IMAC FF columns (GE Healthcare) which were pre-equilibrated with 50mM phosphate buffer with 500 mM NaCl and 20 mM imidazole at pH 7.4. The columns were washed with 10 column volumes (CVs) of 50 mM phosphate buffer with 500 mM NaCl and 20 mM imidazole at pH 7.4 followed by eluted with a linear gradient of 20 mM to 500 mM imidazole over 20 CVs. The purified his-tagged CspZ or CspZ-YA was dialyzed against PBS or TBS and verified with the integrity and purity greater than 90% using SDS-PAGE (**Fig. S3**) before storage at −80°C.

### Generation of CspZ-YA mouse monoclonal and rabbit polyclonal antibodies

Purified his-tagged CspZ-YA was provided to Genemed Synthesis, Inc (San Antonio, TX) to generate CspZ-YA IgG (Total). The IgGs in the immunized rabbits containing CspZ-YA IgGs and naïve rabbit IgGs were purified using protein A chromatography and formulated in 1X PBS with 10% BSA, pH 7.4. To retrieve the CspZ-YA (Total), we first covalently conjugated CspZ onto AminoLink Plus Coupling Resin (ThermoFisher Scientific, Waltham, MA) based on the manufacturer’s instructions. The 5 mL IgGs from CspZ-YA immunized rabbits were subsequently mixed to 100 µL CspZ-conjugated resin slurry (resin to PBST buffer ratio as 1:1) for four hours to capture CspZ-YA IgG (Total). The resin was then washed twice with 1X PBST buffer (1X PBS with 0.05% Tween 20). Finally, the captured CspZ-YA IgG (Total) was eluted with 0.65ml of glycine buffer (0.1 M Glycine HCl at pH 2.5) and dialyzed against 1X PBST buffer.

To isolate the CspZ-YA IgG (FH-binding sites) and CspZ-YA IgG (non-FH binding sites) from CspZ-YA IgG (Total), 100 µL of CspZ-conjugated resin slurry was mixed with excess Factor H (Complement Technology, Tyler, TX, USA) for two hours, allowing the formation of CspZ-FH. This step is essential to block the FH binding site on CspZ. After washing off the excess FH using 1X PBST buffer that CspZ-FH resin was mixed with the CspZ-YA IgG (Total) for 90 minutes. After incubation and the centrifugation of the resin, the unbound fraction containing CspZ-YA IgG (FH-binding sites) was collected from the supernatant. The resulting resin was then washed with 1X PBST, and the bound fraction containing CspZ-YA IgG (non-FH-binding sites) was eluted using 0.1 M glycine HCl at pH 2.5. However, FH was found to be eluted with CspZ-YA IgG (non-FH-binding sites); thus, to remove that FH, the eluted proteins were further dialyzed against 1X PBST, followed by protein A affinity purification to capture the CspZ-YA IgG (non-FH-binding sites). The purified CspZ-YA IgG (non-FH-binding sites) was eventually eluted from protein A resin using the glycine buffer and dialyzed against 1X PBST.

The medium samples from each of the eight clones of mouse hybridoma cells producing CspZ-YA mouse monoclonal antibodies were generated by Protein and Monoclonal Antibody Production Core at Baylor College of Medicine. Each of these medium samples was then applied to NA Protein G Spin Columns to purify the CspZ-YA mouse monoclonal IgG (ThermoFisher Scientific) and then quantitated as described in the vendor’s manual.

### ELISAs

To verify the ability of rabbit polyclonal and mouse monoclonal CspZ-YA IgGs in recognizing FH-binding sites of CspZ, we compared the levels of each of these IgGs in binding to FH-saturated CspZ with those in binding to CspZ proteins treated with BSA (control). One µg of histidine-tagged CspZ was coated on ELISA plate wells, followed by being blocked with 5% BSA in PBS buffer. Those wells were subsequently treated with human FH (500 nM, ComTech, Tayler, TX), BSA (control, 500 nM, Sigma-Aldrich, St. Louis, MO), or PBS (control), followed by incubation with each of the tested rabbit polyclonal and mouse monoclonal CspZ-YA IgGs (50 nM). HRP-conjugated goat anti-mouse IgG (Sigma-Aldrich) and goat anti-rabbit IgG (Sigma-Aldrich) were used to detect the binding of mouse monoclonal and rabbit polyclonal CspZ-YA IgGs, respectively. Tetramethylbenzidine solution (ThermoFisher) was added to each well and incubated for five minutes, then the reaction was stopped with hydrosulfuric acid. Plates were read at 405 nm using a Tecan Sunrise Microplate reader (Tecan, Morrisville, NC). The resulting absorption values were normalized to those from PBS-treated wells to obtain the percentage of CspZ-YA IgG binding.

To determine the ability of rabbit polyclonal and mouse monoclonal CspZ-YA IgGs in preventing FH binding to CspZ, the ELISA was performed as described with modifications (32). Each ELISA microtiter well was coated with one µg of histidine-tagged CspZ. After being blocked with 5% BSA in PBS buffer, the wells were incubated with PBS (control) or serially-diluted irrelevant mouse IgG (anti-green fluorescence protein of mouse IgG, ThermoFisher) or irrelevant rabbit IgG (anti-green fluorescence protein of rabbit IgG, ThermoFisher), or each of the mouse monoclonal or rabbit polyclonal CspZ-YA IgGs (0.4 nM, 0.8 nM, 1.6 nM, 3.125 nM, 6.25 nM, 12.5 nM, 25 nM, 50 nM) followed by being mixed with 500 nM of human FH. Sheep anti-human FH (1:200×, ThermoFisher) and then goat anti-sheep HRP (1:2000×, ThermoFisher) were added, and the levels of FH binding were detected by ELISA as mentioned above. Data were expressed as the proportion of FH binding from serum-treated to PBS-treated wells. The 50% inhibitory concentration (IC_50_) (**Table S1**), representing the IgG concentration that blocks 50% of FH binding, was calculated using dose-response stimulation fitting in GraphPad Prism 5.04.

The seropositivity of the mice after infection with *B. burgdorferi* was determined by detecting the presence or absence of the antibodies that recognize C6 peptides, which has been commonly used for human Lyme disease diagnosis (55). 50 µl of serially diluted mouse serum (1:100×, 1:300×, 1:900×) from 21dpf was added to microtiter wells coated with C6 peptides ((55), Genemed Synthesis, Inc). Total IgG was detected using HRP-conjugated goat anti-mouse IgG (1:20,000×; Bethyl, Montgomery, TX, USA). After the incubation with antibodies for one hour, tetramethyl benzidine solution (ThermoFisher) was added, and the absorbance was detected at 620nm for 10 cycles of 60-second kinetic intervals with 10 seconds shaking duration using Tecan Sunrise Microplate reader as described above. For each serum sample, the maximum slope of optical density/minute of all the dilutions was multiplied by the respective dilution factor, and the greatest value was used as representative of antibody titers (arbitrary unit (A.U.)). The seropositive mice were defined as the mice with the serum samples yielding a value greater than the threshold, the mean plus three-fold standard deviation of the IgG values derived from the uninfected mice.

### Borreliacidal assays

The ability of CspZ-YA mouse monoclonal and rabbit polyclonal IgGs to eradicate spirochetes was determined as described with modifications (32). Briefly, irrelevant mouse or rabbit IgG, or each of these CspZ-YA mouse monoclonal or rabbit polyclonal antibodies were serially diluted to the indicated concentrations (0.4 nM, 0.8 nM, 1.6 nM, 3.125 nM, 6.25 nM, 12.5 nM, 25 nM, 50 nM). The diluted IgGs were then mixed with complement-preserved guinea pig serum (Sigma-Aldrich; final concentration 5%). Note that this concentration of guinea pig sera serum has been verified to not kill spirochetes in the absence of CspZ-YA antibodies (data not shown). The PBS treatment was included as a control. After incubated with the strains B31-5A4, 297, or VS461, the mixture was incubated at 33°C for 24 hours. Surviving spirochetes were quantified by directly counting the motile spirochetes using dark-field microscopy and expressed as the proportion of IgG-treated to PBS-treated Lyme borreliae. The 50% borreliacidal titer is shown in Table S2, representing the serum dilution rate that kills 50% of spirochetes, which was calculated using dose-response stimulation fitting in GraphPad Prism 5.04.

### Generation of infected ticks

Generating infected *I. scapularis* ticks has been described previously (56). BALB/c C3-deficient mice were infected intradermally with 10^5^ of the strains B31-5A4 (27, 48). Ear tissues were collected via ear punch, and bacterial gDNA was purified for detection with qPCR to confirm infection (see section “IgG inoculation, *B. burgdorferi* infection, quantification of spirochete burdens”). Approximately 100 to 200 uninfected larvae were then allowed to feed to repletion on the infected mice as described previously (56). The engorged larvae were collected and allowed to molt into nymphs in a desiccator at room temperature with 95% relative humidity and light-dark control (light to dark, 16:8 hours).

### IgG inoculation, *B. burgdorferi* infection, and quantification of spirochete burdens

Three-week-old, female C3H/HeN mice were subcutaneously inoculated with irrelevant mouse or rabbit IgG or each of the tested CspZ-YA mouse monoclonal or rabbit polyclonal IgGs (1 mg/kg). Five mice per group were used in this study. This number was justified by the power analysis. Using the means ± standard deviation from this study (10±10 for the uninfected control group, 55±10 for the infection group), a power analysis for attaining a statistically significant difference (p <0.05; a one-way ANOVA) between the negative control group and other groups with 95% probability requires minimum of 5 animals per group (57). This value is consistent with numbers that we and others have used in the past for similar studies (32, 58). At 24 hours after inoculation, five nymphs carrying *B. burgdorferi* strain B31-5A4 were allowed to feed to repletion on each mouse, and a subset of nymphs was collected pre- and post-feeding as described (32, 48). Mice were sacrificed at 21 dpf to collect the biting site of skin, knees, heart, and bladder. DNA was purified from these nymphs, and spirochetes were quantified as described with modifications (32). DNA was purified using EZ-10 Spin Column Animal Genomic DNA Mini-Prep Kit (Bio Basic, Inc., Markham, Ontario, CA). Spirochete burdens were quantified based on the amplification of *recA* from *B. burgdorferi* strain B31-5A4 using the primers ((BBRecAfp (5’-GTGGATCTATTGTATTAGATGAGGCTCTCG-3’) and BBRecArp (5’-GCCAAAGTTCTGCAACATTAACACCTAAAG-3’)) with qPCR using an Applied Biosystems 7500 Real-Time PCR system (ThermoFisher) in conjunction with PowerUp™ SYBR® Green Master Mix (ThermoFisher) as described (33, 59). The number of *recA* copies was calculated by establishing a threshold cycle (Cq) standard curve of a known number of *recA* genes extracted from strain B31-5A4, and burdens were reported as the number of *recA* copies per tick or normalized to 100ng of total DNA and reported as the number of *recA* copies per 100ng of total DNA.

### Statistical analyses

Significant differences were determined with a Mann-Whitney test (between two groups)(60), Kruskal-Wallis test with the two-stage step-up method of Benjamini, Krieger, and Yekutieli (61) (more than two groups), or two-tailed Fisher test (for seropositivity in Table S3)(62) using GraphPad Prism 5.04. A p-value < 0.05 was used to determine significance.

## ACKNOWLEDGEMENTS

The authors thank John Leong for providing *B. burgdorferi* strains B31-5A4 and 297, as well as *B. afzelii* strain VS461. They also thank Nicholas Mantis, Klemen Strle, and ChingLin Hsieh for valuable advice. The authors also thank the Wadsworth Animal Core for assistance with Animal Care. The authors appreciate Dean Edwards, Yingmin Zhu, Kurt Christensen, Karen Moberg from Baylor College of Medicine Protein and Monoclonal Antibody Production Core for the consultation and generation of the CspZ-YA monoclonal antibody. This work was supported by NIH grant R21AI144891 (Y.C., A.L.M., T.A.N., M.E.B, W.H.C., Y.L. R.T.K, ZL) and NCI- CA125123 (supporting the generation of CspZ-YA monoclonal antibody in Baylor College of Medicine Protein and Monoclonal Antibody Production Core). The funders had no role in study design, data collection, interpretation, or the decision to submit the work for publication. The authors have no conflict of interest to declare.

## Notes

### Competing Interest Statement

The authors have declared no competing interest.

