## Supplementary information for "CspZ FH-binding sites as epitopes promote antibody-mediated Lyme borreliae clearance"

**Supplemental Table 1: $IC_{50}$ values of the CspZ-YA IgGs tested in this study for blocking FH binding to CspZ from *B. burgdorferi* B31-5A4.**

**$IC_{50}$ (nM)**

| **Rabbit IgG** | | | |
|---|---|---|---|
| **Irr. rab. IgG**[a] | **CspZ-YA IgG (Total)** | **CspZ-YA IgG (Non-FH-binding sites)** | **CspZ-YA IgG (FH-binding sites)** |
| ni.[b] | 19.17±2.87 | ni. | 4.20±0.63 |

| **Mouse IgG** | | | | | | | | |
|---|---|---|---|---|---|---|---|---|
| **Irr. ms. IgG**[c] | **142** | **224** | **582** | **605** | **651** | **1009** | **1139** | **1193** |
| ni. | 82.02±10.12 | ni. | ni. | 57.90±1.85 | 97.69±8.94 | ni. | 4.20±0.72 | 4.07±0.59 |

[a]Irrelevant rabbit IgG, anti-green fluorescence protein of rabbit IgG.

[b]No FH binding inhibition was detected after incubation with 50nM of indicated IgG (the maximal IgG dose used in this study).

[c]Irrelevant mouse IgG, anti-green fluorescence protein of mouse IgG.

**Supplemental Table 2: $BA_{50}$ values of the CspZ-YA IgGs used in this study**

| Mixed with | $BA_{50}$ (nM) Rabbit IgG | | | |
|---|---|---|---|---|
| | Irr. rab. IgG[a] | CspZ-YA IgG (Total) | CspZ-YA IgG (Non-FH-binding sites) | CspZ-YA IgG (FH-binding sites) |
| ***Bb* B31-5A4[b]** | nk.[c] | 7.40±0.20 | 153.30±9.01 | 2.56±0.08 |
| ***Bb* 297[d]** | nk. | 12.85±0.43 | nd.[e] | 3.03±0.31 |
| ***Ba* VS461[f]** | nk. | 7.94±1.07 | nd. | 5.48±0.71 |

| Mixed with | Mouse IgG | | | | | | | | |
|---|---|---|---|---|---|---|---|---|---|
| | Irr. ms. IgG[g] | 142 | 224 | 582 | 605 | 651 | 1009 | 1139 | 1193 |
| ***Bb* B31-5A4** | nk. | 82.02±10.12 | nk. | nk. | 57.90±1.85 | 97.69±8.94 | nk. | 3.45±0.06 | 1.23±0.01 |
| ***Bb* 297** | nk. | nd. | nd. | nd. | nd. | nd. | nd. | 2.48±0.12 | 1.27±0.02 |
| ***Ba* VS461** | nk. | nd. | nd. | nd. | nd. | nd. | nd. | 7.13±0.53 | 51.60±1.92 |

[a]Irrelevant rabbit IgG, anti-green fluorescence protein of rabbit IgG.

[b]*B. burgdorferi* strain B31-5A4

[c]No killing was detected after incubation with 50nM of indicated IgG (the maximal IgG dose used in this study).

[d]*B. burgdorferi* strain 297

[e]Not determined

[f]*B. afzelii* strain VS461

[g]Irrelevant mouse IgG, anti-green fluorescence protein of mouse IgG.

**Supplemental Table 3: Strains and plasmids used in this study.**

| Strain or plasmid | Genotype or characteristic | Source |
|---|---|---|
| *B. burgdorferi* | | |
| B31-5A4 | Clone 5A4 of *B. burgdorferi* B31 isolated from *I. scapularis* ticks in US. | (1) |
| 297 | Clone A11/B11 of *B. burgdorferi* 297 isolated from human Cerebrospinal fluid from US. | (2, 3) |
| *B. afzelii* | | |
| VS461 | Clone JL of *B. afzelii* VS461 isolated from *I. ricinus* ticks in Switzerland. | (4) |
| *E. coli* | | |
| BL21(DE3) | F−, *ompT hsdSB* (rB− mB−) *gal dcm* (DE3) | Novagene |
| BL21(DE3)/pET41a-CspZ | BL21(DE3) producing residues 19 to 237 of CspZ followed by a TEV protease cleavage site and hexa-histidine | This study |
| BL21(DE3)/pET41a-CspZ-YA | BL21(DE3) producing residues 19 to 237 of CspZ-YA followed by a TEV protease cleavage site and hexa-histidine | This study |
| Plasmids | | |
| pET41a-CspZ | KanR[a]; pET41a encoding protein residue 19 to 237 of CspZ followed by a TEV protease cleavage site and hexa-histidine | This study |
| pET41a-CspZ-YA | KanR; pET41a encoding protein residue 19 to 237 of CspZ-YA followed by a TEV protease cleavage site and hexa-histidine | This study |

[a] Kanamycin resistant

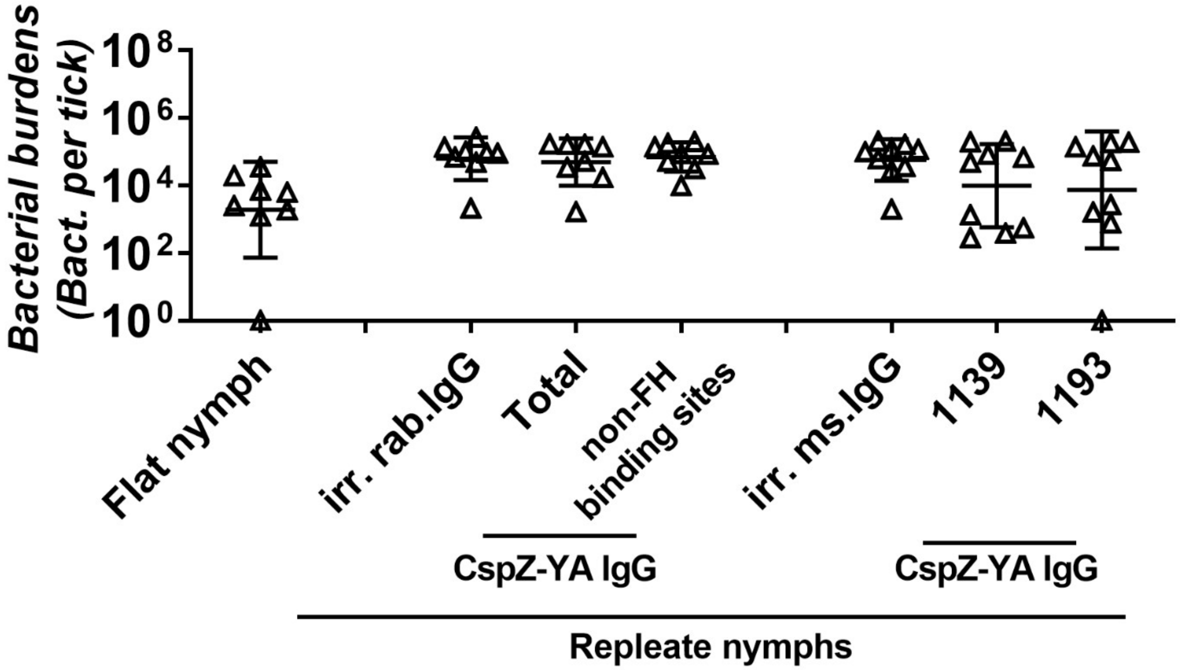


**Supplemental Figure 1. Passive inoculation of CspZ-YA IgGs does not eliminate *B. burgdorferi* B31-5A4 in ticks feeding on mice.** C3H/HeN mice were inoculated with irrelevant IgG from rabbits (irr. rab. IgG) or mice (irr. ms. IgG), or CspZ-YA IgG samples (1mg/kg, five mice per group). These CspZ-YA IgGs include total CspZ-YA IgG (Total), those IgGs that recognize non-FH-binding site (non-FH-binding sites), or mouse monoclonal IgGs #1139 or 1193. At 24 hours after IgG inoculation, these mice were fed on by *I. scapularis* nymphs carrying *B. burgdorferi* B31-5A4 (*Bb* B31-5A4) and those nymphs feeding to repletion were collected. The nymphs prior to feeding were also included as control (Flat nymphs). Spirochete burdens at those nymphs were quantitatively measured and shown as the number of spirochetes per nymph (Bact. per tick). Data shown are the geometric mean ± geometric standard deviation of the bacterial burdens from eight flat nymphs or the nymphs feeding on mice inoculated with irrelevant rabbit IgG, total CspZ-YA IgG, or those IgG that recognize non-FH-binding site, or nine nymphs feeding on mice inoculated with irrelevant mouse IgG or the mouse monoclonal antibody #1139 or 1193.

No statistical significances ($p > 0.05$, Kruskal-Wallis test with the two-stage step-up method of Benjamini, Krieger, and Yekutieli) of differences in bacterial burdens between the groups of nymphs were detected.

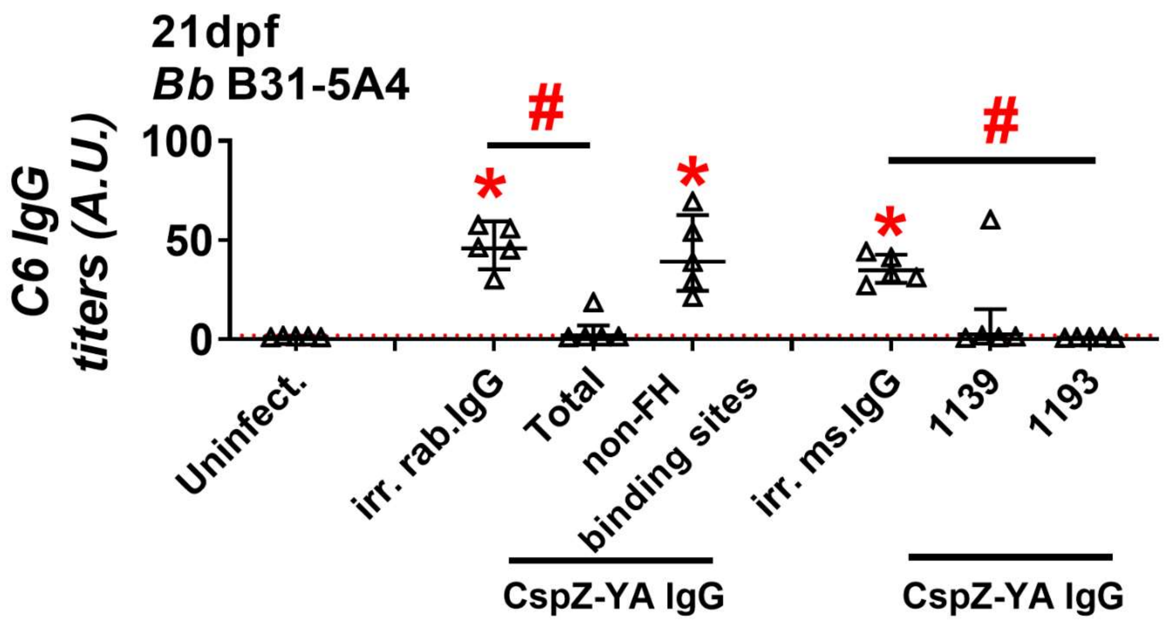


**Supplemental Figure 2. CspZ-YA IgGs that recognize CspZ FH-binding sites selectively prevent seropositivity caused by *B. burgdorferi* B31-5A4 infection.** C3H/HeN mice were inoculated with irr. IgG from rabbits (irr. rab. IgG) or mice (irr. ms. IgG), or CspZ-YA IgG samples (1 mg/kg, five mice per group). These CspZ-YA IgGs include total CspZ-YA IgG (Total), those IgGs that recognize non-FH-binding site (non-FH-binding sites), or mouse monoclonal IgG #1139 or 1193. At 24 hours after IgG inoculation, these mice were fed on by *I. scapularis* nymphs carrying *B. burgdorferi* B31-5A4 (*Bb* B31-5A4). An additional five mice inoculated with PBS but not fed on by ticks were included as the control (Uninfect.). The sera were collected from those mice at 21dpf. Seropositivity was determined by measuring the levels of IgG against C6 peptides in the sera of those mice were using ELISA. The mouse was considered as seropositive if that mouse had IgG levels against C6 peptides greater than the threshold, the mean plus three-fold standard deviation of the IgG levels against C6 peptides from the PBS-inoculated, uninfected mice (red dotted line). The number of mice in each group with the anti-C6 IgG levels greater than the threshold (seropositive) is shown in Table 1. Data shown are the geometric mean ± geometric standard deviation of the titers of anti-C6 IgG. Statistical significances ($p < 0.05$, Kruskal-Wallis test with the two-stage step-up method of Benjamini, Krieger, and Yekutieli) of differences in IgG titers relative to (*) uninfected mice or (#) between indicated groups of mice are presented.

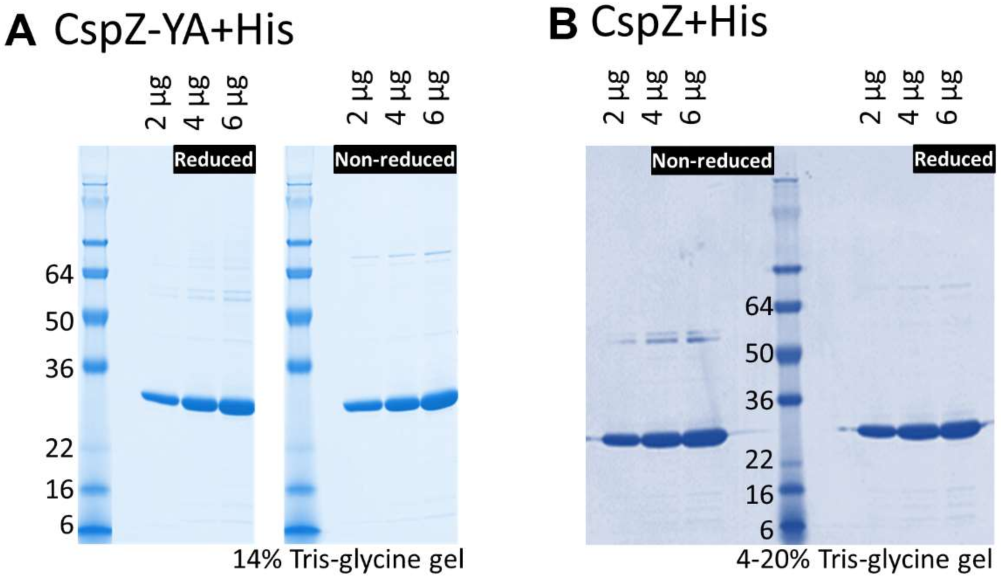


| Loading (μg) | Purity (%) Non-reduced | reduced |
|---|---|---|
| 2 | 93.0 | 93.8 |
| 4 | 93.6 | 94.0 |
| 6 | 93.1 | 93.9 |
| **Mean** | **93.2±0.3** | **93.9±0.1** |

| Loading (μg) | Purity (%) Non-reduced | reduced |
|---|---|---|
| 2 | 92.4 | 94.6 |
| 4 | 90.2 | 95.3 |
| 6 | 93.1 | 95.5 |
| **Mean** | **91.9±1.2** | **95.1±0.4** |

**Supplemental Figure 3. Purity assessment for the purified his-tagged CspZ-YA and CspZ.** Two to six micrograms of CspZ-YA or CspZ were loaded onto 14% or 4-20% tris-glycine SDS-PAGE gels. The purity of each of these proteins was analyzed by densitometry and shown at the bottom panels.
